# Holocentromere diversity in Cyperaceae: contrasting repeat organisation in Mapanioideae and Cyperoideae

**DOI:** 10.64898/2026.06.15.732478

**Authors:** Mariela Sader, Yennifer Mata-Sucre, Yi-Tzu Kuo, Veit Schubert, Thiago Nascimento, Joerg Fuchs, Yhanndra Dias, Klaus Pistrick, Nafiseh Sargheini, Bruno Huettel, Andre Luis Laforga Vanzela, André Marques, Andreas Houben, Andrea Pedrosa-Harand

**Author notes:** Multidisciplinary Institute of Plant Biology (IMBIV), University of Córdoba–CONICET, Córdoba, Argentina (Mariela Sader). Department of Life Sciences, National Cheng Kung University, Tainan City, Taiwan (Yi-Tzu Kuo).

## Abstract

Centromeres ensure accurate chromosome segregation and are typically confined to a single, localised region in monocentric chromosomes. In contrast, holocentric chromosomes exhibit kinetochore activity distributed along the chromosome length. Although holocentricity is widespread in Cyperaceae, the composition and organisation of these centromeres, as well as their evolutionary diversification, remain poorly understood. Here, we investigated centromere organisation in representatives of the subfamilies Mapanioideae (*Hypolytrum schraderianum* Nees) and Cyperoideae (*Cladium mariscus* (L.) Pohl) by combining genome assemblies, repeatome characterisation (RepeatExplorer), fluorescence in situ hybridisation (FISH), and immunolocalisation. Comparative synteny analyses incorporating the genomes of *Rhynchospora breviuscula* (*n* = 5) and *Carex littledalei* (*n* = 29) identified conserved blocks, eventually expanding almost whole chromosomes of *H. schraderianum* (*n* = 30) and *Cl. mariscus* (*n* = 39), despite divergent chromosome numbers and deep evolutionary distances within Cyperaceae. Mobile elements showed very low abundances and were uniformly dispersed, with Ty1/Copia Angela being the most abundant in both species. In *Cl. mariscus*, holocentromeres showed an extended distribution of centromere- and kinetochore-associated proteins along the chromosomes, largely colocalised with two satellite DNA repeats that form dispersed clusters. In contrast, *H. schraderianum* also displayed kinetochore signals along chromatids, but the most abundant satellite DNA family was enriched in distal and interstitial chromosomal regions rather than interspersed along the chromatids. Together, these results reveal different genomic architectures underlying holocentric organisation in phylogenetically distinct Cyperaceae lineages, suggesting that holocentromeres in this family have diversified with variation in centromere organisation in regard to its association with repetitive DNA.

## INTRODUCTION

Centromeres are specialised chromosomal regions that recruit kinetochore proteins and mediate spindle microtubule attachment, ensuring accurate chromosome segregation during mitosis and meiosis (Talbert & Henikoff, 2020). In most eukaryotes, centromeres assemble a conserved kinetochore machinery that connects chromatin to spindle microtubules through the KMN network (KNL1–MIS12–NDC80 complex; Neumann et al., 2023; Oliveira et al., 2024). In classical monocentric species, centromere activity is restricted to a single localised chromosomal domain at the primary constriction. In contrast, holocentric chromosomes lack a primary constriction and exhibit kinetochore activity distributed along most of their length, allowing microtubule attachment at multiple sites (Marques & Drinnenberg, 2025). This organisation enables holocentric chromosomes to better tolerate large-scale chromosomal rearrangements, such as fusions and fissions, because chromosome segregation is not impaired (Hofstatter et al., 2022; Mata-Sucre et al., 2024; Zhang et al., 2026).

Holocentric chromosomes have evolved repeatedly across eukaryotes, including several independent origins in angiosperms (Marques & Drinnenberg, 2025). Although the ancestral centromere organisation in eukaryotes remains debated, monocentricity is generally considered the ancestral state (Schubert et al., 2020; Senaratne et al., 2022). Within flowering plants, holocentric chromosomes are particularly prominent in the Cyperaceae family (Marques et al., 2015; Chung et al., 2018). Holocentricity was historically regarded as a synapomorphy of the Cyperid clade, which includes Cyperaceae, Juncaceae, and Thurniaceae (Greilhuber, 1995; Chase et al., 2006; Givnish et al., 2010), but the discovery of monocentric representatives in related lineages, such as *Juncus* and *Prionium*, has challenged this view (Guerra et al., 2019; Baez et al., 2020). These findings suggest that centromere organisation may be evolutionarily more dynamic than previously assumed and highlight the need to better understand the diversity of centromere architectures. Recent genomic studies have revealed substantial diversity in the structure of centromeres across plant lineages, including satellite-associated holocentromeres in *Rhynchospora* (Marques et al., 2015) and *Chionographis* (Kuo et al., 2023), non-satellite-based holocentromeres *in Luzula elegans* (Heckmann et al., 2013) and *Myristica fragrans* (Kuo et al., 2024), CENH3-independent holocentromeres in *Cuscuta europaea* (Oliveira et al., 2020), macro-monocentromeres in *Chamaelirium luteum* (Kuo et al., 2025). and meta-polycentromeres in *Pisum* and *Lathyrus* species (Neumann, et al., 2016). These findings highlight the need for comparative analyses across phylogenetically informative taxa to better understand the origin and diversification of holocentric chromosomes in Cyperaceae.

Cyperaceae comprises approximately 5,500 species divided into two major subfamilies, Mapanioideae and Cyperoideae, and exhibits remarkable karyotypic diversity (Roalson, 2008; Semmouri et al., 2019). Chromosome numbers vary widely within the family, particularly in Cyperoideae, whereas the few investigated representatives of Mapanioideae displayed comparatively lower variability (Márquez-Corro et al., 2018; Dias et al., 2020). Despite holocentric chromosome structure being well studied in core Cyperoideae, including *Rhynchospora* and *Eleocharis* (Marques et al., 2015; Ribeiro et al., 2017; Souza et al., 2024), the genomic architecture underlying holocentric chromosomes in other Cyperaceae lineages remains poorly understood. In particular, the genomic architecture underlying holocentric chromosomes and the distribution of kinetochore-associated components in these taxa remain largely unknown. To address this gap, we selected *Hypolytrum schraderianum* and *Cladium mariscus*, which represent phylogenetically distinct lineages of the two major Cyperaceae subfamilies, Mapanioideae and Cyperoideae, respectively. While *H. schraderianum* is a perennial geophyte that grows primarily in wet tropical biomes across South and Tropical America, *Cl. mariscus* is a perennial wetland plant with an almost cosmopolitan distribution, naturally occuring across temperate, tropical, and subtropical regions on every continent except Antarctica. These species provide an opportunity to investigate whether key features of holocentric chromosome organisation have been maintained across deep evolutionary timescales within the family.

Building on this phylogenetic framework, we investigated centromere organisation and repeat composition in *Hypolytrum schraderianum* (Mapanioideae and *Cladium mariscus* (deeply divergent Cyperaceae lineages from subfamily Cyperoideae). We integrated genome assemblies, repeatome characterisation, fluorescence *in situ* hybridisation (FISH) of repeats, and immunolocalisation of the kinetochore component KNL1 (Oliveira et al., 2024; Neumann et al., 2023), α-tubulin fibers, and the cell cycle-dependent pericentromeric phosphorylation of histone H3 serine 28 (H3S28ph) mark (Goto et al., 2002; Gernand et al., 2003), to compare genomic features associated with holocentric chromosomes in these taxa. This comparative approach allowed us to assess whether key features of holocentric chromosome organisation are shared between these phylogenetically distinct representatives of Cyperaceae or whether distinct centromere landscapes are associated with their contrasting genomic architectures. We hypothesised that key cytological hallmarks of holocentric chromosomes would be conserved in both species despite their deep evolutionary divergence, whereas centromere sequence organisation could exhibit lineage-specific diversification

## MATERIALS AND METHODS

### Plant materials

Individuals of *Hypolytrum schraderianum* Nees were collected in the Brazilian Atlantic Forest, while *Cladium mariscus* (L.) Pohl was collected in Seeland, Germany. All plants were cultivated under greenhouse conditions at the Leibniz Institute of Plant Genetics and Crop Plant Research (IPK), Gatersleben, Seeland, and at the Max Planck Institute for Plant Breeding Research, Cologne, Germany. Detailed collection information is provided in Table 1.

**Table 1.** Species of Cyperaceae used for genome sequencing and genome size estimations, collection sites, and vouchers.

| Species | 1C<br>(pg/Mbp) | 2n | Collection sites and vouchers | GenBank<br>accession |
| --- | --- | --- | --- | --- |
| *Cladium mariscus* | 0.283/277 | 78 | Deutschland, Sachsen-Anhalt, Gatersleben (GAT 115 651), ca. 2.3 km southwest of Schwanebeck, leg. K. Pistrick 27.7.2022. | XXXX |
| *Hypolytrum schraderianum* | 0.340/332 | 60 | Brazil, Paraná: Ilha do Mel, Paranaguá; voucher FUEL 29197 (Dias, M.C.; Rocha, E., s.n., 2 July 1995), barcode FUEL019443; coordinates: -25.52, -48.509201 (WGS84; ±24,916 m) | XXXX |

### Genome size estimation

The nuclear DNA content of *H. schraderianum* and *Cl. mariscus* was estimated by flow cytometry following Dolezel et al. (2007), with minor modifications. Nuclear suspensions were prepared by co-chopping ∼0.5 cm² of fresh young leaf tissue together with *Raphanus sativus* L. (2C = 1.11 pg) as an internal reference standard in a Petri dish using the CyStain PI Absolute P reagent kit (Sysmex-Partec), according to the manufacturer’s instructions. The suspension was filtered through a 50 µm CellTrics mesh (Sysmex-Partec) and analysed using a CyFlow Space flow cytometer (Sysmex-Partec, Germany). Three individuals per species were analysed, with two independent replicates per individual. The absolute nuclear DNA content (2C, pg) was calculated as the ratio between the mean fluorescence of the sample and that of the internal standard multiplied by the known 2C value of the reference standard.

### PacBio Sequencing

High-molecular-weight DNA was isolated from 1,5-gram fresh weight material with the Nucleobond HMW kit (Macherey Nagel). Quality was assessed by capillary electrophoresis (Agilent FEMTOpulse), and DNA was quantified with Qubit HS (Fisher Scientific). Next, DNA was directly used as input for a barcoded PacBio SPK3.0 library as recommended by the vendor. Then, an additional size selection was done on BluePippin (Sage Science) to enrich for DNA fragments larger than 8 kb. The recovered library was sequenced on a Sequel IIe at the Max Planck Genome-centre Cologne for 30 hours, and high-fidelity (HiFi) data was post-run generated.

### Hi-C library

For Hi-C library preparation, nuclei were extracted with the Sigma Cell TMPN plant nuclei isolation/extraction kit. The Hi-C library was prepared with the Arima High Coverage HiC Kit according to the user guide for plant tissues, and the library was sequenced with Illumina NextSeq 2000 in 2 x 150 bp paired-end read mode.

### Genome assembly and Hi-C scaffolding

HiFi and Hi-C reads were assembled using Hifiasm v0.16.1 (Cheng et al., 2021) with the following command: hifiasm -o output.asm -t 40 <u>reads.fq.gz</u>. Assemblies were evaluated for contiguity and completeness using BUSCO (Seppey et al., 2019) and QUAST (Gurevich et al., 2013). Hi-C reads were mapped to the primary contigs generated by Hifiasm using BWA (Li & Durbin, 2009) following the hic-pipeline workflow (https://github.com/esrice/hic-pipeline).

Hi-C–based scaffolding was performed with SALSA2 (Ghurye et al., 2019) using default parameters and specifying the restriction sites “GATC, GAATC, GATTC, GAGTC, GACTC”. Several minimum mapping quality (MAPQ) thresholds were tested, and the final scaffolding was performed using MAPQ ≥ 10. Following automated scaffolding, multiple rounds of manual curation were conducted based on visual inspection of Hi-C contact heatmaps. Regions displaying inconsistent or multiple contact patterns were manually reorganised using Juicebox (Durand et al., 2016) and the 3D-DNA assembly pipeline (Dudchenko et al., 2017) to correct scaffold position and orientation and generate pseudomolecules.

### Genome annotation

Gene prediction was performed using the Helixer web server (Stiehler et al., 2021) with default parameters and the genome assembly masked by RepeatMasker v4.1.5 (Smit et al., 2015) as input. Transposable elements (TE) and tandem repeats were annotated using the DANTE, DANTE_LTR, and TideCluster tools from the RepeatExplorer2 pipeline (https://repeatexplorer-elixir.cerit-sc.cz/) using the complete genome assemblies as input (Novák et al., 2024). All putative tandem sequences were compared for homology using DOTTER (Sonnhammer & Durbin, 1995) and individually mapped to the genome using BLAST with 95% similarity in Geneious. The resulting TE and tandem repeat annotation files were converted to BED format and used as input tracks for genome visualization with ShinyCircos (Yu et al., 2018) and pyGenomeTracks using 100 kb windows. Interstitial telomeric sequences (ITSs) were annotated at two stringency levels by identifying regions longer than 200 bp with ≥75% or ≥90% similarity to the *Arabidopsis*-type telomeric repeat (TTTAGGG) using Geneious (Kearse et al., 2012). Arrays located within 10 kb were merged to avoid over-fragmentation of closely spaced telomeric repeats, which likely represent a single interstitial telomeric locus separated by small insertions, sequence divergence, or assembly gaps. This approach provides a more accurate estimate of ITSs abundance and genomic distribution. Representative satellite DNA arrays were further analysed using ModDotPlot (Nadalin et al., 2024) to visualise internal repeat structure and sequence homogenisation. Self-alignment dot plots and sequence identity distributions were used to compare the organisation of satellite arrays between species.

For the *in-silico* analysis of the repetitive fraction in the assembled genomes, repeat annotation was performed using CARP (Comprehensive Annotation of Repeats Pipeline; Kavonrtep, 2025), following the developer’s official protocol available on GitHub (https://github.com/kavonrtep/CARP). This Snakemake-based workflow enables the identification, classification, and quantification of repetitive sequences in genome assemblies. The pipeline integrates DANTE (Novák et al., 2024) for transposable element annotation and classification, and TideCluster (https://github.com/kavonrtep/TideCluster) for the detection and characterisation of tandem repeats, allowing the identification of both dispersed and tandem repetitive elements.

### LTR retrotransposon identification and phylogenetic analysis

Complete retrotransposon nucleotide sequences and reverse transcriptase (RT) amino acid sequences were manually extracted from DANTE-LTR output files using Linux-based grep commands. Both datasets were partitioned by superfamily and subsequently re-annotated using the NCBI Conserved Domain Database (CDD) (Wang et al., 2023). Only sequences containing all essential domains in the correct order and exhibiting consistent inter-domain distances were considered intact, applying a coverage threshold of >80%. Graphical representations were generated using IBS-Illustrator (Xie et al., 2022). Phylogenetic reconstruction was performed based on RT amino acid sequences, retaining only those with >80% coverage and sequence identity. Sequences were aligned using ClustalW, and a maximum likelihood tree was inferred with 500 bootstrap replicates under an appropriate amino acid substitution model. The resulting tree was visualised using iTOL v7.2.1 (Letunic and Bork, 2024).

### Synteny analysis

To place our analyses in a broader evolutionary context, we compared *H. schraderianum* and *Cl. mariscus* genomes with other well-characterised holocentric species from core Cyperoideae (*Rhynchospora breviuscula* and *Carex littledalei*) for which high-quality genome assemblies are available (GenBank accessions: *Rhynchospora breviuscula*, GCA_027562975, and *Ca. littledalei*, GCA_011114355). Synteny relationships were inferred using the DEEPSPACE pipeline based on pairwise whole-genome alignments. Genome assemblies were aligned in pairs, and collinear blocks were identified and visualised as a synteny map linking homologous regions across chromosomes. Divergence times between lineages were incorporated to provide evolutionary context, using estimates of ∼82.6 million years ago (Mya) for the split between *Hypolytrum* (Mapanioideae) and Cyperoideae (represented here by *Cladium*, *Rhynchospora*, and *Carex*), ∼74.4 Mya for the divergence between *Cladium* and the lineage leading to *Rhynchospora* and *Carex*, and ∼69.6 Mya for the divergence between *Rhynchospora* and *Carex* (Escudero et al., 2013).

### Satellite DNA probe generation and fluorescence in situ hybridisation (FISH)

Following the identification and characterization of satellite DNA families (see Results), primers for the most abundant satellite DNA families HscSAT1-181 and HscSat2-184 satDNAs from *H. schraderianum* were designed oriented outwards using Primer3 (Untergasser et al., 2012) implemented in Geneious v9.1 (Table S2). PCR reactions (50 μL) contained 20 ng genomic DNA, 0.1 mM dNTPs, 2 mM MgClC, 1×PCR buffer, 0.4 μM of each primer, 0.4×TBT, and Taq DNA polymerase. PCR amplification consisted of 30 cycles of 95°C for 1 min, annealing for 1 min, and 72°C for 1 min. PCR products were sequenced using an ABI 3500 sequencer (Applied Biosystems) to confirm amplification of the satDNA motifs, and probes were labeled by nick translation with Atto488 or Atto550 (Jena Bioscience).

Consensus sequences of the most abundant putative satellite repeats of *Cl. maricus* were used to design oligonucleotide probes (Table S2). Fluorochrome-conjugated oligos were synthesised by Eurofins (Germany) and used to detect the satellite repeats CmaSat1-92 (TAM) and CmaSat2-172 (FAM). The clone pAtT4 (Richards and Ausubel, 1988) was used as a probe to detect the *Arabidopsis*-type telomere repeat. Plasmid DNA from this clone was labeled with ATTO488-dUTP using a Nick Translation Labeling Kit (Jena Bioscience, Germany).

For slide preparations, roots were pretreated with 2mM 8-hydroxyquinoline for 24 h at 4°C, fixed in ethanol:acetic acid (3:1, v/v) for 2 h, and stored at −20°C. After enzymatic digestion with 2% cellulase, 2% pectinase, 2% pectolyase in citrate buffer (0.01 M sodium citrate dihydrate and 0.01 M citric acid) for 2 h 30 min at 37°C, mitotic slides were prepared by air-drying following Carvalho & Saraiva (1993), with modifications according to Ribeiro et al. (2020). FISH was performed as described by Aliyeva-Schnorr et al. (2015), and the slides were counterstained with 2 µg/ml DAPI (4′,6-diamidino-2-phenylindole) (Sigma) in antifade (Vectashield, Vector).

### Immunodetection

Mitotic chromosomes and interphase nuclei were prepared from root tips fixed in 4% paraformaldehyde with 1% Igepal in Tris buffer (10 mM Tris, 10 mM EDTA, 100 mM NaCl, 0.1% Triton X-100, pH 7.5), washed, and enzymatically digested at 37 °C for 45–60 min using a mixture containing 2% cellulase, 2% pectolyase, and 2% cytohelicase in 1× PBS buffer. Meristems were chopped in LB01 nuclei isolation buffer, filtered through a 50-µm CellTrics filter, and centrifuged onto Superfrost Plus slides using a Cytospin3 centrifuge. Primary antibodies diluted in 2% BSA in 1×PBS containing 0.5% Triton X-100 and 0.2% Igepal were applied and incubated at 37 °C for 1 h, followed by overnight incubation at 4 °C. After washing, slides were incubated with secondary antibodies, dehydrated through an ethanol series, air-dried, and counterstained with DAPI in antifade medium (Vectashield). Primary antibodies included rabbit anti-*Cuscuta europaea* KNL1 (1:400) (Oliveira et al., 2024), rat anti-histone H3S28ph (1:1000) (Sigma-Aldrich, cat. no. H9908), and mouse anti-α-tubulin (1:300) (Sigma-Aldrich, cat. no. T9026-2). For microtubule immunodetection, the pretreatment was omitted, and buffers were replaced with microtubule-stabilising buffer (MTSB). The immunodetection procedure followed previously described protocols with minor modifications (Oliveira et al., 2024).

Immuno-FISH was performed following Dias et al. (2024). After immunostaining and imaging, coverslips were removed, and the DAPI-containing mounting medium was washed off with 1×PBS. Slides were post-fixed in freshly prepared 3:1 ethanol:glacial acetic acid for 10 min at room temperature and air-dried in darkness. Before hybridisation, slides were pre-incubated overnight at 37 °C in a humid chamber with the hybridisation mixture containing probes. Slides were then washed in 2×SSC for 5 min and dehydrated through an ethanol series (70%, 90%, and 100%; 3 min each). DNA denaturation was performed in 0.2 N NaOH in 70% ethanol for 10 min at room temperature, followed by a brief wash in ice-cold 1×PBS, dehydration in an ethanol series (70%, 90%, 100%), and air-drying. Probe denaturation, hybridisation, and counterstaining were performed as described above for FISH, except that the stringent wash was carried out at room temperature.

### Microscopy and image analysis

Chromatin ultrastructure and signal localisation were analysed via spatial structured illumination microscopy (3D-SIM, super-resolution) on an Elyra PS.1 system applying the ZENBlack software (Carl Zeiss GmbH) (Weisshart et al., 2016). The 3D-SIM image stacks were used to generate animations with the Imaris 9.7 (Bitplane) software.

## RESULTS

### Genome size and assemblies

The genome size of *H. schraderianum* estimated by flow cytometry was 2C = 0.68 pg (Table 1), corresponding to ∼332 Mb for the haploid 1C genome. The assembled genome of *H. schraderianum* (2*n* = 60) yielded a total size of 276.2 Mb (276,199,152 bp) long, with 30 scaffolds corresponding to chromosome-scale pseudomolecules and 148 smaller unplaced contigs. The scaffold N50 was ∼8 Mb, whereas the contig N50 was ∼306 kb (Table S1). BUSCO analysis revealed 98.4% complete single-copy orthologs (87.3% single-copy and 11.1% duplicated), with 1.4% fragmented and 0.2% missing BUSCOs (n = 425), indicating a highly complete genome assembly (Fig. S1a-c; Fig. 1a).

**Figure 1.**
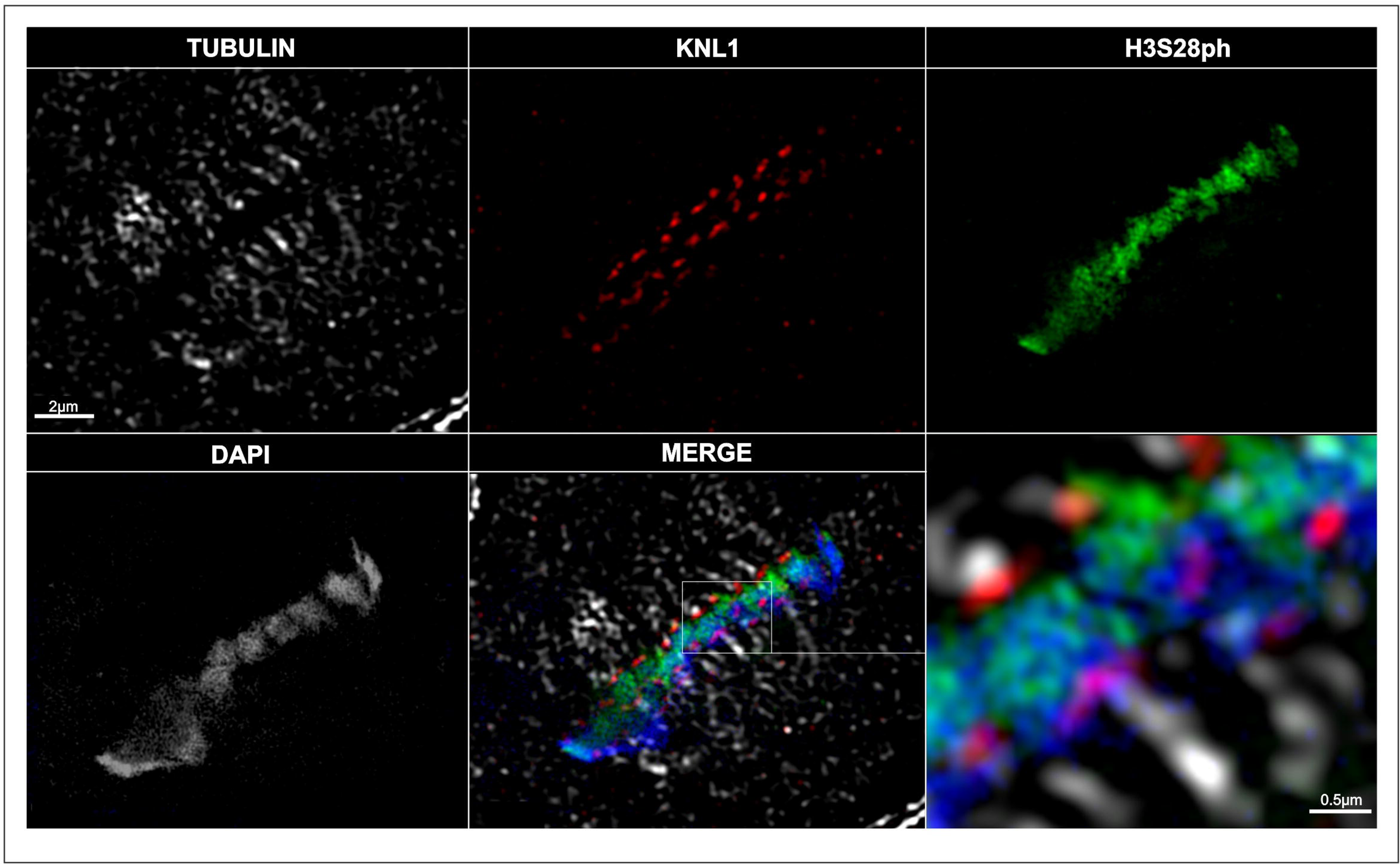
Circos plots showing the chromosomal distribution of genes and major repetitive sequence families in *H. schraderianum* (a) and *Cl. mariscus* (b). (a) From the outer to the inner tracks: gene density, satellite DNA families HscSAT1_181 and HscSAT2_184, followed by Ty1/Copia and Ty3/Gypsy LTR retrotransposons. (b) From the outer to the inner tracks: gene density (blue), satellite repeats (CmaSat1-92, CmaSat2-172, CmaSat3-434), and transposable elements (Ty1/Copia, Ty3/Gypsy) across chromosomes. Histograms represent gene density calculated using a 100 kb window size. The outer heatmap indicates sequence density, where blue represents lower abundance and red represents higher abundance. c) ModPlot self-alignment and genomic distribution of HscSat1-181 and HscSat2-184 in a representative region of chromosome 1 of *Hypolytrum schraderianum*. The boxed region in the annotation tracks corresponds to the genomic interval analyzed by ModPlot. d) ModPlot analysis of a representative CmaSat1-92 region in chromosome 1 of *Cladium mariscus*. The highlighted interval corresponds to the region analysed in the self-alignment plot.

The genome size of *Cl. mariscus* estimated by flow cytometry was 2C = 0.566 pg (Table 1), corresponding to ∼277 Mb for the haploid 1C genome. The genome assembly of *Cl. mariscus* (2*n* = 78) yielded a total size of 303.7 Mb (303,663,834 bp). The assembled genome slightly exceeded the size estimated by flow cytometry, what may reflect residual redundancy in the assembly, such as incompletely collapsed haplotypic regions or duplicated repetitive sequences. The assembly included 39 chromosome-scale scaffolds (pseudomolecules) and 1,423 smaller unplaced contigs, with N50 values of ∼6 Mb and ∼34.3 kb, respectively (Table S1). BUSCO completeness reached 99.5% (97.2% single-copy and 2.4% duplicated), with 0.5% fragmented and no missing BUSCOs (n = 425), also indicating a highly complete assembly (Fig. S1d-f; Fig. 1b). Overall, both assemblies exhibit high completeness and contiguity with a low proportion of unresolved bases (10.46 Ns per 100 kbp in *H. schraderianum* and 3.76 Ns per 100 kbp in *Cl. mariscus*), corresponding to a total of 26.5 kb and 9.5 kb of ambiguous sequence, respectively. GC content distributions were unimodal in both species, with around ∼35% GC, indicating a relatively homogeneous nucleotide composition and no evidence of distinct compositional compartments (Table S1; Fig. S1).

Chromosome size distributions inferred from the assemblies revealed differences between species. *Hypolytrum schraderianum* exhibited larger chromosomes overall (median ≈ 8.3 Mb; range ≈ 5.5–12 Mb), whereas *Cl. mariscus* showed smaller chromosomes (median ≈ 6.0 Mb; range ≈ 4.2–11 Mb). In both species, chromosome sizes were relatively evenly distributed, with no clear evidence of differentiated size classes (Fig. S2).

### Repeat composition and organization

#### Hypolytrum schraderianum

Satellite DNA analysis identified four satDNA families. The most abundant was HscSat1-181 (Tandem Repeat Cluster, TRC_1), representing 2.4% of the genome, followed by HscSat2-184 (1.4%), composed of two closely related variants (TRC_2 and TRC_5) (Fig S3) with monomer lengths of 182 and 184 bp. Two additional low-abundance families were detected, representing 0.03% (HscSat3-49) and 0.02% (HscSat4-2603) of the genome. Overall, the identified high-copy satellite DNA families comprised ∼3.85% of the haploid genome (Table S3). Consensus sequences of the satellite DNAs are provided in Table S4.

Satellite array size analysis revealed marked differences among satDNA families (Fig. S4). HscSat1-181 exhibited a broad distribution of array sizes (median of 5.7 Kb), including several large, clustered arrays (up to 0.33 Mb), whereas HscSat2-184 was generally restricted to smaller, scattered arrays with lower variability (median of 3.3 Kb up to 58.9 Kb; Fig. 1c). The two low-abundance satellite families were represented by relatively small and less variable arrays (data not shown). ModPlot analyses in *H. schraderianum* revealed different structural configurations of the Hsc satellite families. In the region chr1:9.20-9.40, for instance, HscSat1-181 forms a long array located adjacent to a short HscSat2-184 array. The corresponding self-alignment plot reveals discrete homogenised blocks (around 90% identity), representing separate tandem arrays (Fig. 1c). Overall, these patterns indicate differential amplification dynamics among satellite families within the genome.

The validated full-length LTR retrotransposons identified by DANTE-LTR accounted for 13.97 Mb, corresponding to only 4.20% of the haploid genome. Most elements belonged to the Ty1/Copia superfamily, which comprised 11.37 Mb (3.42% of the genome), whereas Ty3/Gypsy elements represented 2.60 Mb (0.78%). Within Ty1/Copia, Angela was the most abundant lineage (1.95% of the genome), followed by Ale (0.72%), Ikeros (0.46%), Tork (0.18%), Ivana (0.06%), TAR (0.05%), and SIRE (0.01%). Among Ty3/Gypsy elements, Tekay was the most abundant lineage (0.28% of the genome), followed by Athila (0.24%), Ogre (0.19%), and minor contributions from Galadriel and Reina (each ∼0.04%) (Table S5).

The *in-silico* distribution of Ty1/Copia and Ty3/Gypsy elements in the *H. schraderianum* genome revealed a relatively even distribution across all chromosomes, with no clear evidence of large-scale compartmentalisation (Fig. 1). Most LTR retrotransposon lineages exhibited low to moderate abundance and a dispersed distribution pattern, with no lineage forming extensive continuous domains, similar to the short HscSat2-184 arrays. Among them, Ty1/Copia Angela elements were relatively more abundant, while Ty3/gypsy lineages such as Tekay were present at lower densities. HscSat1-181 tandem repeats, on the other hand, showed one to four peaks, usually one or two, along multiple chromosomes, indicating localised enrichment, while the two low-abundance satellite families are enriched in a single chromosome each. Overall, the repetitive landscape of *H. schraderianum* reflects a combination of dispersed retrotransposons and less abundant satDNAs and locally enriched HscSat1-181, with satellite DNA contributing substantially to the genomic repeat fraction (Fig. S5 and S6).

Phylogenetic analysis based on reverse transcriptase (RT) amino acid sequences revealed well-supported clustering of LTR retrotransposon lineages, consistent with their classification obtained using DANTE-LTR. The maximum likelihood tree showed grouping of Ty3/Gypsys Chromovirus-related elements, such as CRM and Tekay, supporting the robustness of lineage assignment. In *H. schraderianum*, a limited number of CRM elements were identified, with only a few intact copies, including two nearly identical elements located on different chromosomes. Similarly, Tekay elements were represented by a small number of copies (∼15), many of which were highly similar to each other, indicating low sequence divergence and suggesting recent activity or limited diversification (Fig. S7).

#### Cladium mariscus

Satellite DNA analysis identified three major satDNA families. The most abundant was CmaSat1-92, representing approximately 0.52% of the haploid genome, followed by CmaSat2-172 (0.49%) and a low-abundance family, CmaSat3-434 (0.05%) (Table S6). Overall, satellite DNA accounted for ∼1.06% of the haploid genome. Satellite array size differed among satDNA families (Fig. S8 and S9). CmaSat1-92 exhibited the largest array sizes (median of 25.1 Kb) and the widest distribution, including numerous large arrays (up to 0.29 Mb), whereas CmaSat2-172 was generally associated with smaller arrays (median of 10.1 Kb), although occasional larger outliers were observed (up to 0.30 Mb; Fig. S4). ModPlot analyses revealed short tandem arrays characterised by discontinuous triangular self-alignment plots, with variable, but slightly lower homogenization (∼85%; Fig. 1d). Overall, CmaSat1-92 showed a higher tendency for array expansion than CmaSat2-172.

Analysis of validated full-length LTR retrotransposons identified using DANTE-LTR revealed a total of 4.68 Mb, representing only 1.69% of the haploid genome (Table S7). Elements of the Ty1/Copia superfamily were the most abundant, representing 1.38% of the genome (3.82 Mb). Within this group, the Angela lineage was the most represented (0.36%), followed by Tork (0.33%), Ale (0.32%), Ivana (0.15%), and Ikeros (0.13%). Less abundant lineages included TAR (0.07%), Alesia (0.02%), and Bianca (0.01%). In contrast, Ty3/Gypsy elements accounted for a smaller fraction of the genome (0.31%, 0.86 Mb). Among these, Athila was the most abundant lineage (0.14%), followed by CRM (0.11%) and Reina (0.04%), whereas Galadriel (0.01%) and Tekay (0.01%) were present at very low frequencies. Overall, these results indicate that intact LTR retrotransposons constitute a relatively small fraction of the *Cl. mariscus* genome, with a predominance of Ty1/Copia lineages, particularly Angela, Tork, and Ale (Table S7).

*In silico* analysis revealed that repetitive elements are broadly distributed across all *Cladium mariscus* scaffolds, with no clear evidence of compartmentalisation along chromosomes. Tandem repeats represented one of the most prominent components, showing multiple peaks along most scaffolds, whereas Ty1/Copia and Ty3/Gypsy elements were present at lower densities and appeared more evenly dispersed throughout the genome (Fig. 1). At the lineage level, most LTR retrotransposon families exhibited low abundance and a scattered distribution (e.g., Angela and Athila), with no single lineage dominating large chromosomal regions. Minor enrichment patterns were observed for some lineages (e.g., CRM, Ivana and TAR), although these signals remained weak and discontinuous across scaffolds. Overall, the repetitive landscape of *Cl. mariscus* is characterised by a low abundance of intact LTR retrotransposons and a comparatively higher contribution of tandem repeats, supporting a genome organisation in which satellite DNA constitutes a major repetitive component (Fig. S8 and S9).

In *Cladium mariscus*, Ty3/Gypsy Chromovirus elements account for less than 0.2% of the genome (Table S8). Only a few complete Reina elements were detected, some of which contained insertions disrupting conserved domains. Overall, the low abundance and limited diversity of intact Chromovirus elements contrast with the prevalence of satellite DNA, supporting the idea that satellite repeats may play a more prominent role in centromere organisation in this genome (Fig. S9).

The length distribution of LTR retrotransposons varied among repeat lineages in both species and was broadly similar among shared lineages, with Athila and Angela representing the largest elements and several Ty1/Copia lineages, including Ale and Ivana, being generally shorter. *Hypolytrum schraderianum* exhibited a wider range of element lengths and a greater diversity of detected lineages, including additional Gypsy groups such as Ogre and Tekay, which were absent or only weakly represented in *Cl. mariscus* (Fig. S10). In *H. schraderianum*, full-length elements ranged from approximately 4 to 27 kb, while the corresponding 5′ LTR regions were generally between ∼400 and 2,500 bp, with a few longer outliers. Ty3/Gypsy elements, particularly those belonging to the Ogre (∼15–16 kb), Tekay (∼9–10 kb), and Athila (∼10 kb) lineages, tended to exhibit larger full-length sizes compared with Ty1/Copia elements (Angela: ∼8–9 kb; Ale: ∼5 kb), which generally showed shorter element lengths (Fig. S10a). In *Cl. mariscus*, Athila and Angela exhibited the largest median element sizes, generally approaching 10–12 kb, whereas Ale, Ivana, Tork, and Alesia comprised shorter elements, typically ranging between 4 and 6 kb. CRM, TAR, and Ikeros displayed intermediate sizes (Fig. S10b). In both species, elements containing both PBS and TSD signals showed slightly reduced size variability compared to the full dataset, with more constrained size ranges, suggesting that structurally complete elements tend to exhibit more conserved lengths.

### Synteny analysis

Synteny analysis revealed several chromosomal rearrangements among the four species, with chromosome number varying from *n* = 5 to *n* = 39, despite their relatively similar genome sizes (∼252–370 Mb). Comparisons between *H. schraderianum* and *Cl. mariscus* (representing Mapanioideae and Cyperoideae, respectively; divergence ∼82.6 Mya) show alterations in most chromosomes.Nevertheless, chromosome-size syntenic blocks can be identified in several cases, such as between *H. schraderianum* chr. 21 and *Cl. mariscus* chr. 25, and many chromosomes correspond to a large extent to one or two chromosomes of the other species. (Fig. 2). Similarly, synteny between *Cl. mariscus* and *R. breviuscula* (∼74.4 Mya) and *Ca. littledalei* (divergence of ∼69.6 Mya from *Rhynchospora*) is highly fragmented. Mulitple (ca. 6-8) *Cl. mariscus* chromosomes correspond to each *R. breviuscula* chromosome, while usually 1-3 *Cl. mariscus* chromosomes or chromosome segments corresponded to *Ca. littledalei* chromosomes, with exceptions. Overall, these patterns indicate that large synteny blocks were fused or fissed, reshaping the karyotypes of these Cyperaceae lineages.

**Figure 2.**
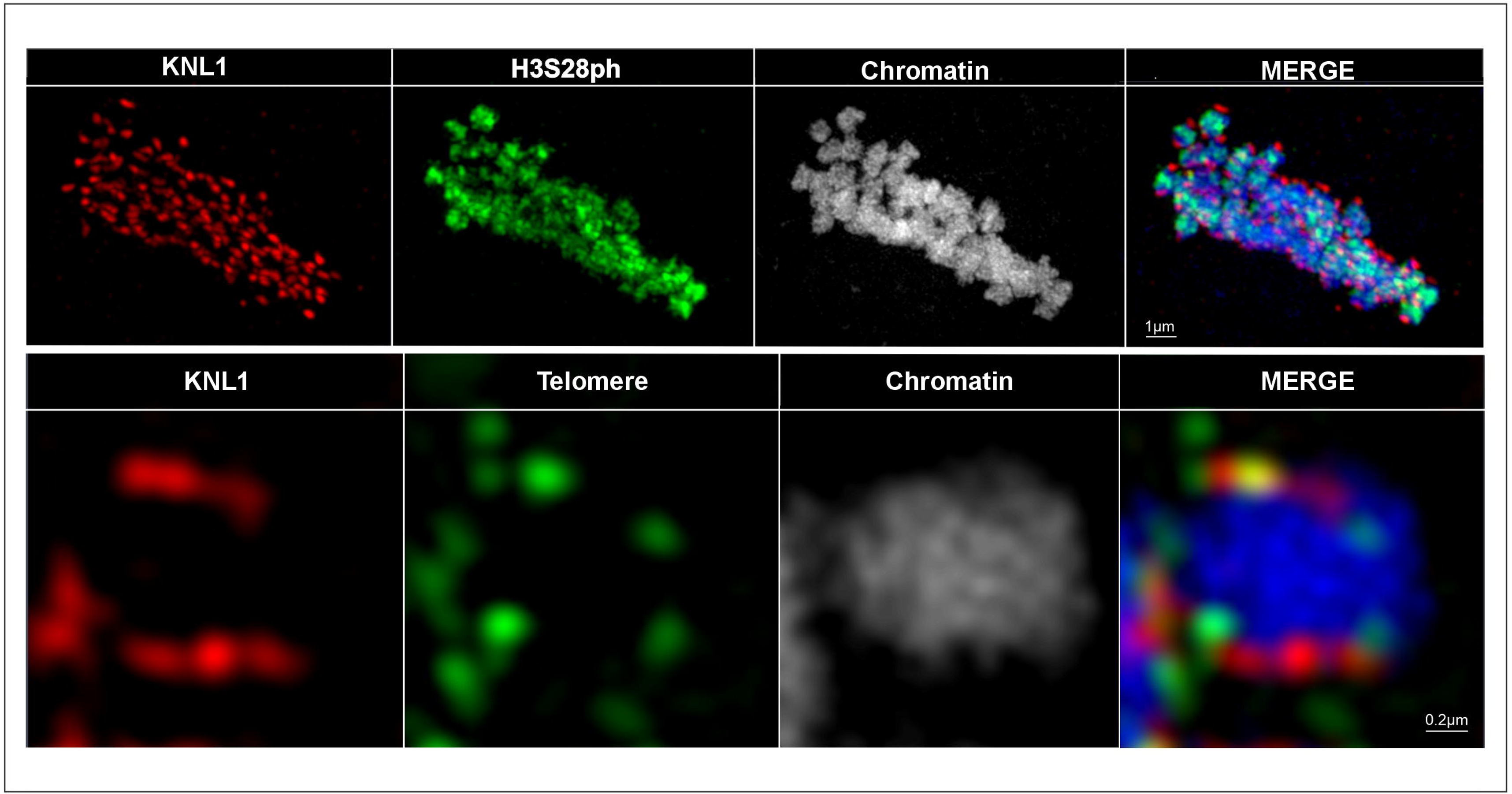
DEEPSPACE synteny map of four holocentric Cyperaceae species. Colored links indicate collinear regions between chromosomes scaled by physical size. Extensive interchromosomal connections reveal highly fragmented synteny and pervasive genome rearrangements across lineages diverged ∼70–83 Mya (Escudero et al., 2013).

### Holocentromeric organisation revealed by immunostaining

#### Hypolytrum schraderianum

To investigate centromere organisation, we performed immunolabelling using antibodies against the conserved outer kinetochore protein KNL1, the cell cycle–dependent (peri)centromere-enriched histone H3 serine 28 phosphorylation (H3S28ph), and α-tubulin. At metaphase, H3S28ph signals were distributed along the entire length of the chromosomes and were positioned between the pole-oriented KNL1 signals, which extended along the chromatids rather than being present in a single localised domain. This dispersed distribution is consistent with a holocentric chromosome organisation, in which kinetochore-associated proteins are present along most of the chromosome length. The α-tubulin labeling revealed spindle microtubules surrounding the chromosomes and in proximity to KNL1-positive regions along the chromatids, consistent with chromosome-wide microtubule attachment (Fig. 3).

**Figure 3.**
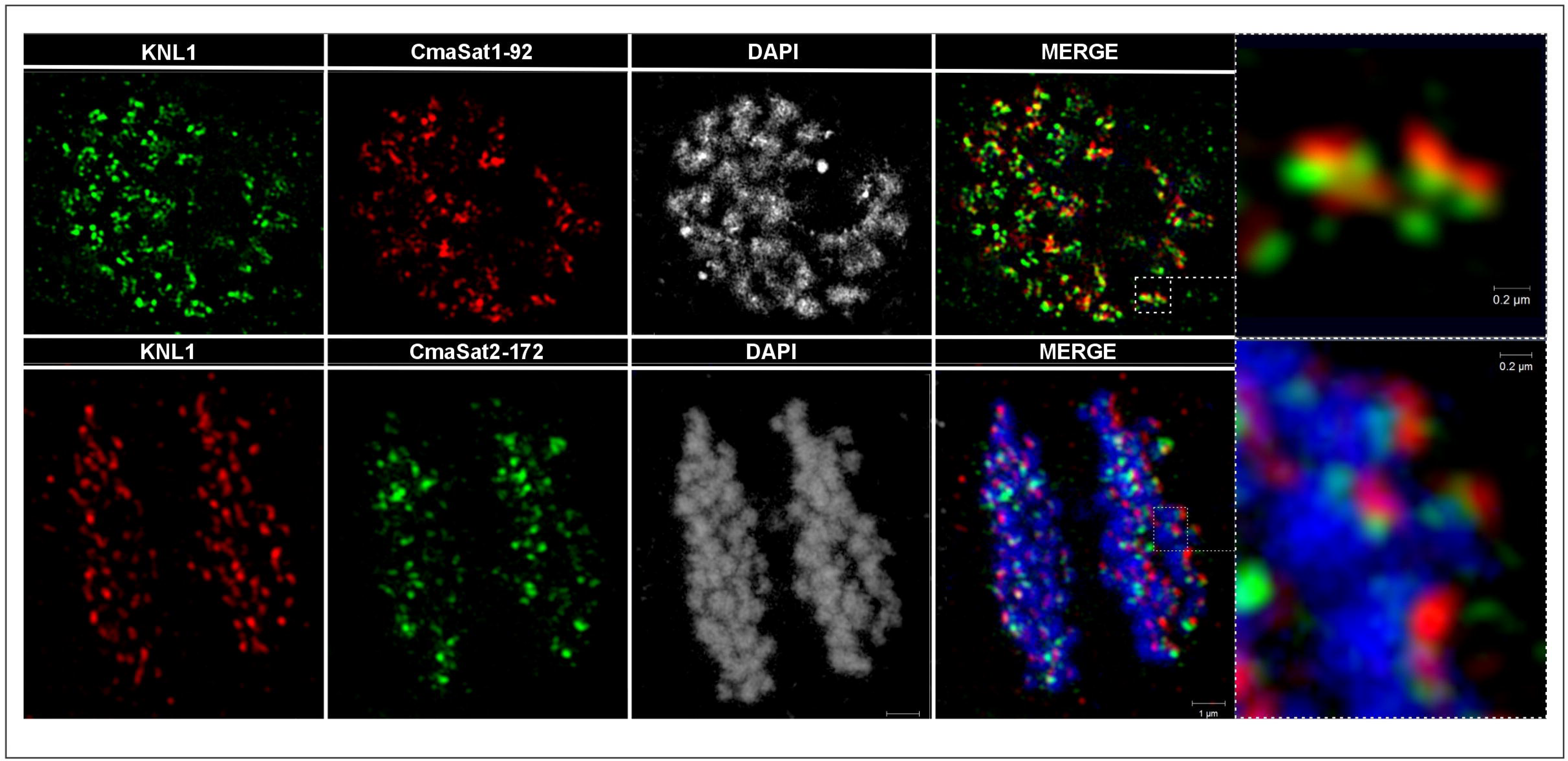
Immunolocalisation of centromere-associated proteins in a somatic *H. schraderianum* metaphase cell. Immunostaining of spindle microtubules (tubulin), KNL1, and phosphorylated histone H3S28. DAPI indicates the whole chromatin. The inset shows a magnified region highlighting KNL1 signals on the outer surface along the chromatids associated with microtubules.

FISH was performed using probes targeting the two major satellite families, HscSat1-181 and HscSat2-184. Neither satellite displayed continuous line-like signals along chromatids that are typically associated with repeat-based holocentromeres (Fig. S11). Instead, probes revealed distinct hybridisation patterns, with HscSat1-181 signals enriched in interstitial (internal) and terminal chromosomal regions and HscSat2-184 more uniformly dispersed, and only partial scolocalisation was observed in some chromosome pairs (Fig. S11).

#### Cladium mariscus

In *Cl. mariscus*, indirect immunostaining using an antibody against KNL1 also revealed an extended distribution of KNL1 signals along the chromatids, indicating a holocentric organisation (Fig. 4; Movie S1-3). However, CmaSat1-92 also showed a continuous distribution along chromatids at metaphase, extending eventually from telomere to telomere. In contrast, CmaSat2-172 displayed a combination of continuous signals along chromatids and discrete dot-like signals located in regions not labelled by CmaSat1-92 (Fig. S12).

**Figure 4.**
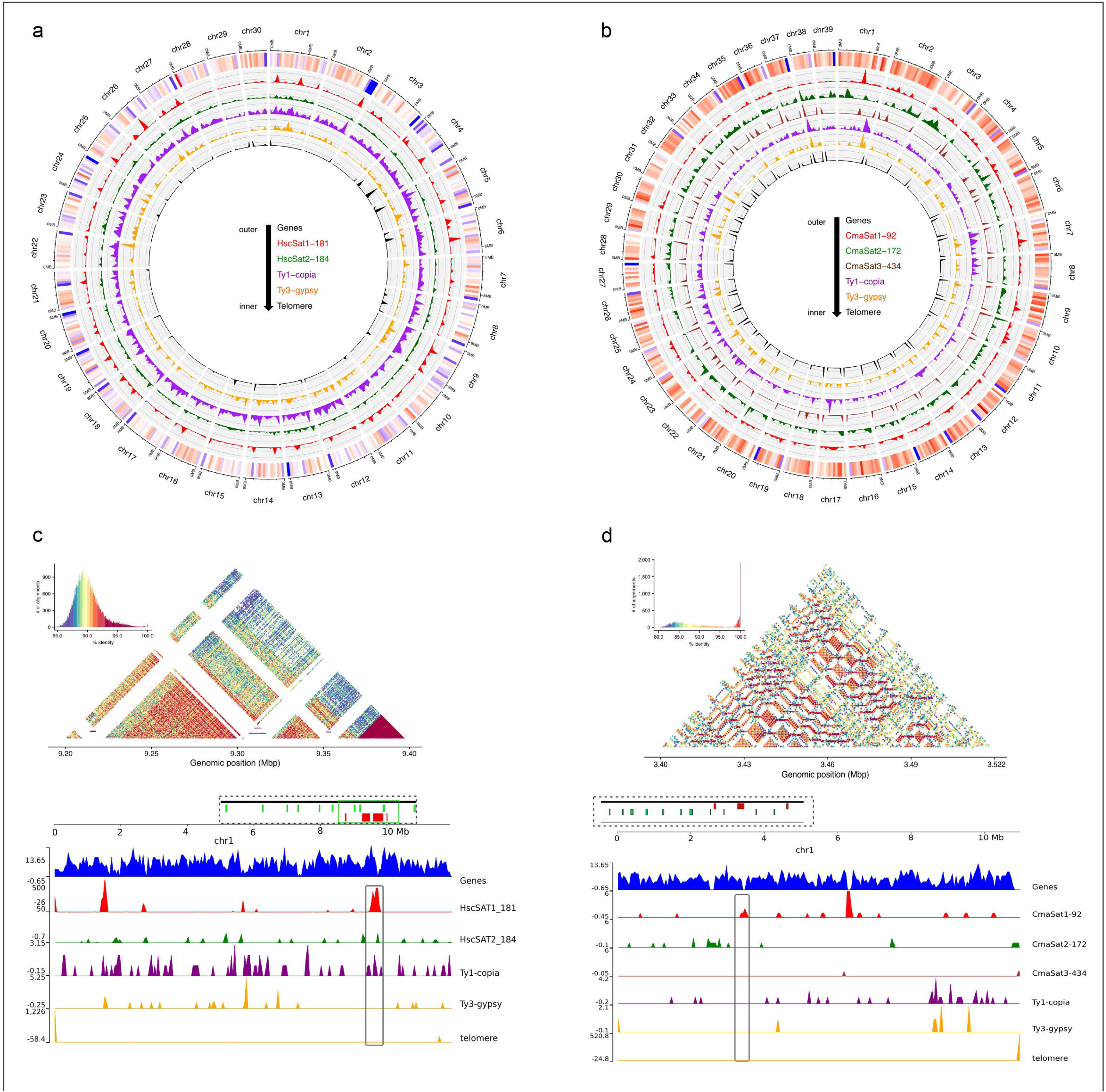
Localisation of centromere-associated proteins and telomeric signals in *Cl. mariscus* somatic metaphase chromosomes. The rectangle in the upper panel indicates the magnified chromosome of the lower panel.

To assess the spatial relationship between satellite DNA and kinetochore distribution, immuno-FISH was performed by combining KNL1 detection with satellite and telomeric probes. KNL1 signals extensively colocalised with CmaSat1-92 along the chromatids, and less frequently with CmaSat2-172, which exhibited a markedly lower degree of association, detected on some chromosomes and in interphase nuclei (Fig. 5; Fig. S13). No specific association was observed with telomeric regions (Figs. 4; Movie S2). This spatial pattern indicates that kinetochore-associated chromatin is enriched in CmaSat1-92 arrays, extending along the chromosome length. CmaSat1-92 defined the principal repeat-rich domains underlying holocentromere organisation in *Cl. mariscus* (Fig. 5).

**Figure 5.**
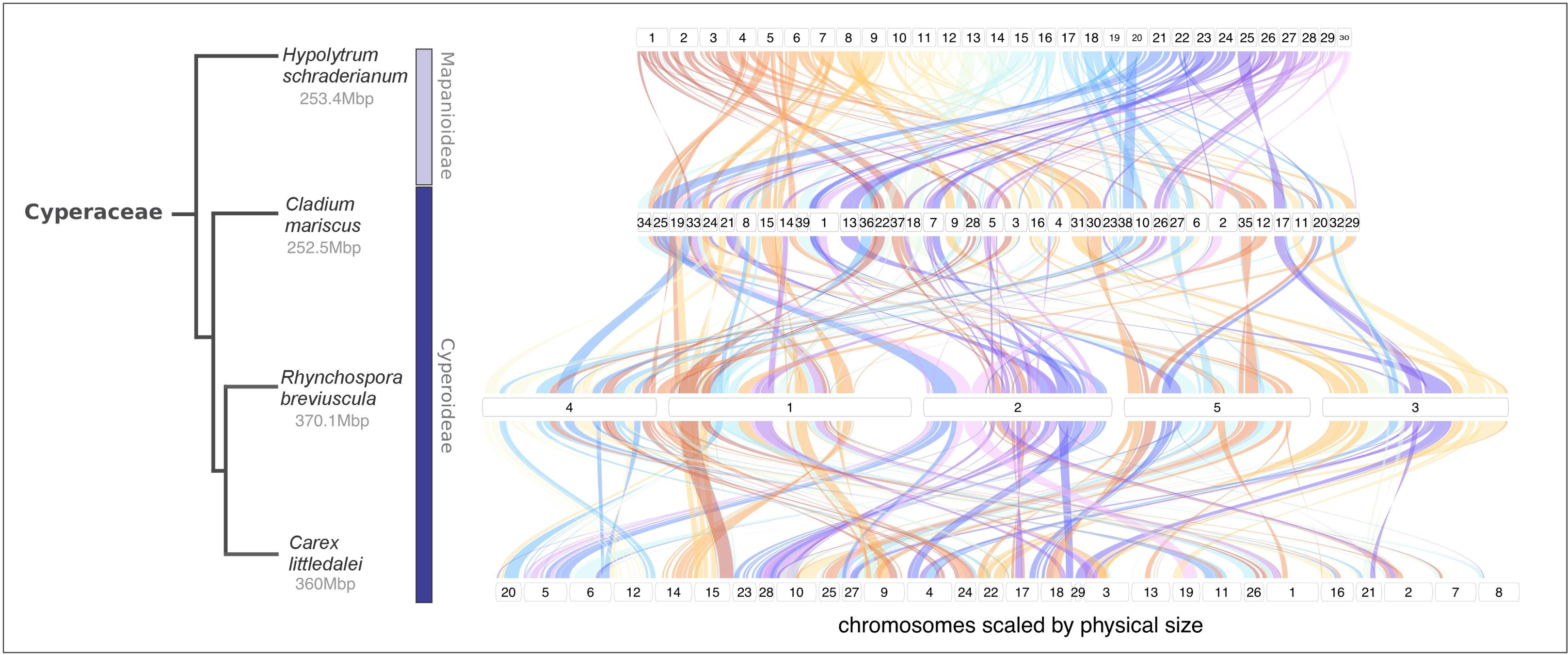
Colocalisation between KNL1 and satellite DNA families in *Cl. mariscus* mitotic chromosomes. Immunofluorescence labeling of KNL1 followed by FISH with probes targeting the satellite repeats CmaSat1-92 and CmaSat2-172. Top row: KNL1, CmaSat1-92, and DNA counterstained with DAPI. The merged image reveals colocalisation between KNL1 and CmaSat1-92 along the chromatids. The inset shows a magnified region highlighting signal overlap. Bottom row: Metaphase chromosomes showing KNL1, CmaSat2-172, and DAPI. The merged image indicates a partial association between KNL1 and CmaSat2-172 signals. The insets show an enlarged region.

## DISCUSSION

### Holocentric chromosome organisation and contrasting kinetochore configurations

By combining genome-scale analyses of repetitive DNA with cytogenetic localisation of centromere-associated proteins in two phylogenetically distant representatives of Cyperaceae, we show that both *H. schraderianum* and *Cl. mariscus* exhibit a kinetochore distribution consistent with holocentric chromosome organisation (Oliveira et al., 2024). However, these species differ in the genomic features associated with centromere organization, revealing distinct structural configurations underlying these holocentric chromosomes.

In *Cl. mariscus*, kinetochore domains were closely associated with extended satellite DNA arrays distributed along the chromosomes, with KNL1 signals extensively colocalising with CmaSat1-92. This pattern indicates that CmaSat1-92 constitutes a major component of the kinetochore-associated chromatin, although it is unlikely to represent the sole determinant of centromere function. In contrast, in *H. schraderianum*, KNL1 signals occurred along chromatids while the major satellite DNA family showed more localised chromosomal distribution. Together, these findings indicate that holocentric chromosomes in these phylogenetically distinct Cyperaceae species can be supported either by satellite DNA-associated or satellite DNA-independent centromere configurations, as described in *Luzula* (Juncaceae) (Mata-Sucre et al., 2024). Despite this difference in major satDNA distribution between both species, further experiments, such as ChIPseq with anti-CENH3, is necessary to determine if less abundant, dispersed satDNA families or mobile elements are enriched in the kinetochore-associated chromatin.

### Repetitive DNA landscapes

Despite their relatively small genome sizes, both species harbour diverse transposable element repertoires, as reported in other plants with compact genomes such as *Arabidopsis thaliana*, *Utricularia gibba*, and *Prionium serratum* (The Arabidopsis Genome Initiative, 2000; Ibarra-Laclette et al., 2013; Baez et al., 2020). In both species, elements of the Ty1/Copia superfamily predominate over Ty3/Gypsy elements. Together, these observations are consistent with the view that genome size is influenced more strongly by transposable element abundance and amplification dynamics (changes in copy number and genomic abundance over evolutionary time) than by transposable element diversity alone (Elliott & Gregory, 2015).

Intact LTR retrotransposons represent a relatively small fraction of the genome in both taxa, although their abundance is slightly higher in *H. schraderianum* than in *Cl. mariscus*. Several lineages are shared, including Angela (Ty1/Copia) and Athila (Ty3/Gypsy). Overall, while both species exhibit a broad distribution of repetitive elements along chromosomes, *Cl. mariscus* is characterized by reduced abundance and diversity of intact LTR retrotransposons and a stronger contribution of tandem repeats, whereas *H. schraderianum* retains a more detectable, albeit moderate, representation of dispersed retrotransposon lineages alongside locally enriched satellite DNA arrays.

### Satellite DNA and its differential role in holocentromere structure

Satellite DNA represents a key component of plant repetitive DNA and often contributes to centromere organisation (Ávila Robledillo et al., 2018; Ribeiro et al., 2017). In *H. schraderianum*, however, themajor satellite families displayed predominantly clustered localisation in terminal and interstitial regions or dispersed distribution. The absence of continuous chromatid-wide signals in metaphase chromosomes suggests that these repeats are not directly associated with kinetochore domains. However, these observations do not exclude the possibility that other repetitive sequences contribute to holocentromere organisation in *H. schraderianum.* This contrasts with the larger and more evenly distributed arrays observed for CmaSat1-92 in *Cl. mariscus*, suggesting distinct evolutionary dynamics of satellite DNA amplification and homogenisation in the two species. Similar diversity in the association between satellite DNA and holocentromeres has been reported among *Rhynchospora* species (Ribeiro et al., 2017).

### Evolutionary implications of holocentromere diversity in Cyperaceae

The species analysed in this study possess relatively small genomes distributed in several small chromosomes. In Mapanioideae, sampling is very limited, but genome sizes range from ∼269 to 367 Mb with chromosome numbers between 2*n* = 46 and 2*n* = 76 (Márquez-Corro et al., 2018; Dias et al., 2020), whereas Cyperoideae exhibits a much wider variation, from 196 Mb to 9,643 Mb and chromosome counts from 2*n* = 4 to 2*n* = 224 (Nishikawa et al., 1984; Kaur et al., 2012; Lipnerova et al., 2013; Ribeiro et al., 2018; Escudero et al., 2014). This variability has often been associated with the holocentric nature of chromosomes, which allows chromosome fragments to retain centromeric activity and facilitates rapid karyotype evolution (Jankowska et al., 2015; Kandul et al., 2007). The observation of holocentromeres in species from diverse lineages within both subfamilies of Cyperaceae supports this hypothesis.

The genome reshuffling observed across *Hypolytrum*, *Cladium*, *Rhynchospora* and *Carex* highlights the dynamic nature of genome evolution in Cyperaceae during ∼75–83 million years. Such reorganisation is consistent with the holocentric chromosome architecture, which has been proposed evolve mostly by multiple chromosomal fissions and fusions, since they lack the constraints imposed by localised centromeres. The dispersed nature of kinetochore activity may allow chromosomal fragments to be stably inherited, promoting karyotype evolution over long evolutionary timescales. In this context, the lack of large, conserved centromeric domains and the presence of small, dispersed satellite arrays further support a model in which both centromere organisation and genome structure are highly flexible in these species.

The genome assemblies and synteny analyses of these phylogenetically distant lineages allow inferences concerning karyotype evolution in the family. The ancestral chromosome number of Cyperaceae has been postulated to be *x* = 5 (Roalson, 2008), with fissions and fusions the major drivers of chromosome evolution in the family. No WGD was inferred for the origin of the family (Winterfeld et al., 2025). A similar hypothesis is considered for the evolution of *Rhynchospora*, one of its most studied genera, but polyploidy is recurrent in several of its lineages (Ribeiro et al., 2018; Burchardt et al., 2020; Hofstatter etal., 2022). Considering the *x* = 5 hypothesis and our present data, after transition to holocentricity, multiple chromosomal fissions occurred in parallel in these different lineages, with numerous small chromosomes believed to increase genetic diversity in this lineage where crossovers are limited during meiosis (Elliott et al., 2022). Polyploidy occurred more recently, largely limited to *Rhynchospora*, being rare in *Carex* (Escudero et al. 2023), and absent in *Cladium* and *Hypolytrum*. Conserved synteny could be the result of conserved genomic hotspots of chromosome rearrangements related to repetitive DNA (Hofstatter etal., 2022; Escudero et al., 2024) and/or limited fusion/fission cycles after chromosome number reduction. Indeed, fusions are more common after polyploidy in *Rhynchospora*, promoting descending dysploidy (Hofstatter et al., 2022). Alternatively, ancestral chromosome number in Cyperaceae was high (*x* = ∼30) before the advent of holocentricity and despite the absence of polyploidy. Under this scenario, the high chromosome number observed in different lineages would be ancestral, explaining the relative conservation of macrosynteny in ∼80 million years. This interpretation is also consistent with the relatively high chromosome numbers reported in the sister lineages of Cyperaceae, *Juncus* and *Prionium* (Báez et al., 2020; Mata-Sucre et al. 2023). Under either scenario, the origin of holocentric chromosomes in Cyperaceae would differ from that proposed for *Luzula*, where holocentric chromosomes have been suggested to originate through the fusion of atypical monocentric chromosomes (Lsy). But the impact of holocentricity would be heterogeneious in this case, more pronounced in some lineages with fewer chromosomes. More whole genome assemblies from different Cyperaceae lineages will help to corroborate one of this hypothesis.

Small monoploid genomes and small chromosomes are likely ancestral in the order Poales, but the evolution of Poaceae went a different pattern. In this family, polyploidy is frequent and also happened before its diversification (Winterfeld et al., 2025). The proposed ancestral grass karyotype (AGK) based on 5 to 7 chromosomes had 12 chromosome pairs after ρ WGD, similar to extant *Oryza sativa* (rice, 2*n* = 2*x* = 24), but descending dysploidy by fusions and insertions mediated by translocations led to lower chromosome number in different lineages, such as the derived *n* = 5 from *Brachypodium distachyon* (Murat et al., 2010), as well as accumulation of repeats leading to very large genomes as observed in *Avena* (Liu *et al*., 2023). The contrasting patterns observed between these related families suggest that the advent to holocentricity was determinant in Cyperaceae, favoring the *x* = 5 followed by fissions’ hypothesis.

Comparative studies in a limited number of holocentric plant species have revealed substantial diversity in centromere organisation, particularly in holocentromere structures (Kuo et al., 2024). For example, *Chionographis japonica* exhibits 7-11 megabase-sized satellite DNA centromeric units (Kuo et al., 2023), whereas *Luzula sylvatica* and *Rhynchospora pubera* possess a high number of short satellite arrays containing centromere unit colocalising with CENH3-containing chromatin (Kuo et al., 2024; Mata-Sucre et al., 2024). Our results place *Cl. mariscus* closer to the latter configuration, with multiple, short satDNA arrays, while *H. schraderianum* represents a distinct condition in which kinetochore activity does not appear to be associated with major repeat arrays. The observed variation highlights the diversity of holocentromere architectures across plant lineages and indicates that different genomic configurations can support holocentric chromosome function. However, the evolutionary relationships among these configurations remain unclear and will require broader taxonomic sampling and functional analyses to be fully resolved.

## Conclusions

Our results reveal diversity in the genomic architecture underlying holocentric chromosomes within Cyperaceae. Although both *H. schraderianum* and *Cl. mariscus* exhibit extended kinetochore activity, their underlying repeat landscapes differ. In *Cl. mariscus*, kinetochore-associated chromatin is closely associated with multiple satellite DNA arrays, whereas in *H. schraderianum*, kinetochore signals were not detectably associated with the major satellite families based on their chromosomal distribution. These findings suggest that holocentric chromosomes in Cyperaceae do not rely on a single conserved structural organisation but may instead be supported by different genomic configurations, including both high-copy repeat-associated and high-copy repeat-independent chromatin contexts.

## Acknowledgments

The authors are grateful to CAPES (Coordenação de Aperfeiçoamento de Pessoal de Nível Superior) for funding through the PROBRAL CAPES/DAAD project (grant number 88881.144086/2017-01) and to CONICET (Consejo Nacional de Investigaciones Científicas y técnicas) for a fellowship awarded to MS. We acknowledge the ELIXIR-CZ Research Infrastructure Project (LM2015047) for providing computational resources for the RepeatExplorer analysis.

## Author Contributions

MS, APH, and AH conceived the study. MS performed the experiments and analysed the data. MS and YMS wrote the original draft of the manuscript. MS, YMS, ALLV, and TN performed bioinformatic analyses. YD and YTK contributed to data analysis and laboratory experiments. VS conducted super-resolution microscopy analyses. JF performed flow cytometry analyses. ALLV and KP collected and identified the plant material. AM, BH, and NS performed the genome assemblies. AH, AM, and APH supervised the study and revised the manuscript. All authors reviewed and approved the final version of the manuscript.

## Supporting Information

Supplementary Figures S1–S15, Tables S1–S8, and Movies S1–S3 supporting this article are available online.

## Data Accessibility

The assembled genomes generated in this study have been deposited in the NCBI database under BioProject accession number [XXXX]. All accession numbers are provided in Supplementary Table S1.

